# Targeting individual cells by barcode in pooled sequence libraries

**DOI:** 10.1101/178681

**Authors:** Navpreet Ranu, Alexandra-Chloé Villani, Nir Hacohen, Paul C. Blainey

## Abstract

There is rising interest in applying single-cell transcriptome analysis and other single-cell sequencing methods to resolve differences between cells. Pooled processing of thousands of single cells is now routinely practiced by introducing cell-specific DNA barcodes early in cell processing protocols^1-5^. However, researchers must sequence a large number of cells to sample rare subpopulations^6-8^, even when fluorescence-activated cell sorting (FACS) is used to pre-enrich rare cell populations. Here, a new molecular enrichment method is used in conjunction with FACS enrichment to enable efficient sampling of rare dendritic cell (DC) populations, including the recently identified AXL^+^SIGLEC6^+^ (AS DCs) subset^7^, within a 10X Genomics single-cell RNA-Seq library. DC populations collectively represent 1-2% of total peripheral blood mononuclear cells (PBMC), with AS DC representing only 1-3% of human blood DCs and 0.01-0.06% of total PBMCs.

---

Here we introduce a simple PCR-based approach to enrich a pooled single-cell library for specific cells of interest based on initial analysis of a shallow sequence dataset (*e.g.*, 5000 reads per cell for RNA-Seq)^9,10^. Researchers can then enrich the library to focus sequencing effort on specific cells of interest. We carry out enrichment using PCR primers specific to the barcodes of target cells to preferentially amplify cognate molecules from the pooled sequence library in multiplexed PCR reactions (10-plex; Fig. 1A, Supplementary Table 1). The resulting libraries are enriched approximately 100-fold for the group of targeted cells and can be sequenced to achieve deep coverage of high-quality target cells at far lower overall sequencing effort than would have been required in sequencing the original library (Fig. 1B, 1C, Supplementary Fig. 1). To test the method, we targeted 24 human primary single cells (selected to represent both low and high quality cells) within a library of 2397 Lineage^−^HLA-DR^+^ cells enriched from PBMCs. RNA abundances determined by sequencing the enriched libraries quantitatively recapitulated RNA abundances from the original library, which was deeply sequenced and computationally resampled to provide matched control datasets for comparison (Fig. 1D, Supplementary Fig. 2-4). Data from the enriched library also resulted in congruent cluster assignments in reduced-dimensional data representations (Fig. 1E, Supplementary Figs. 5 and 6).

**Fig 1.**
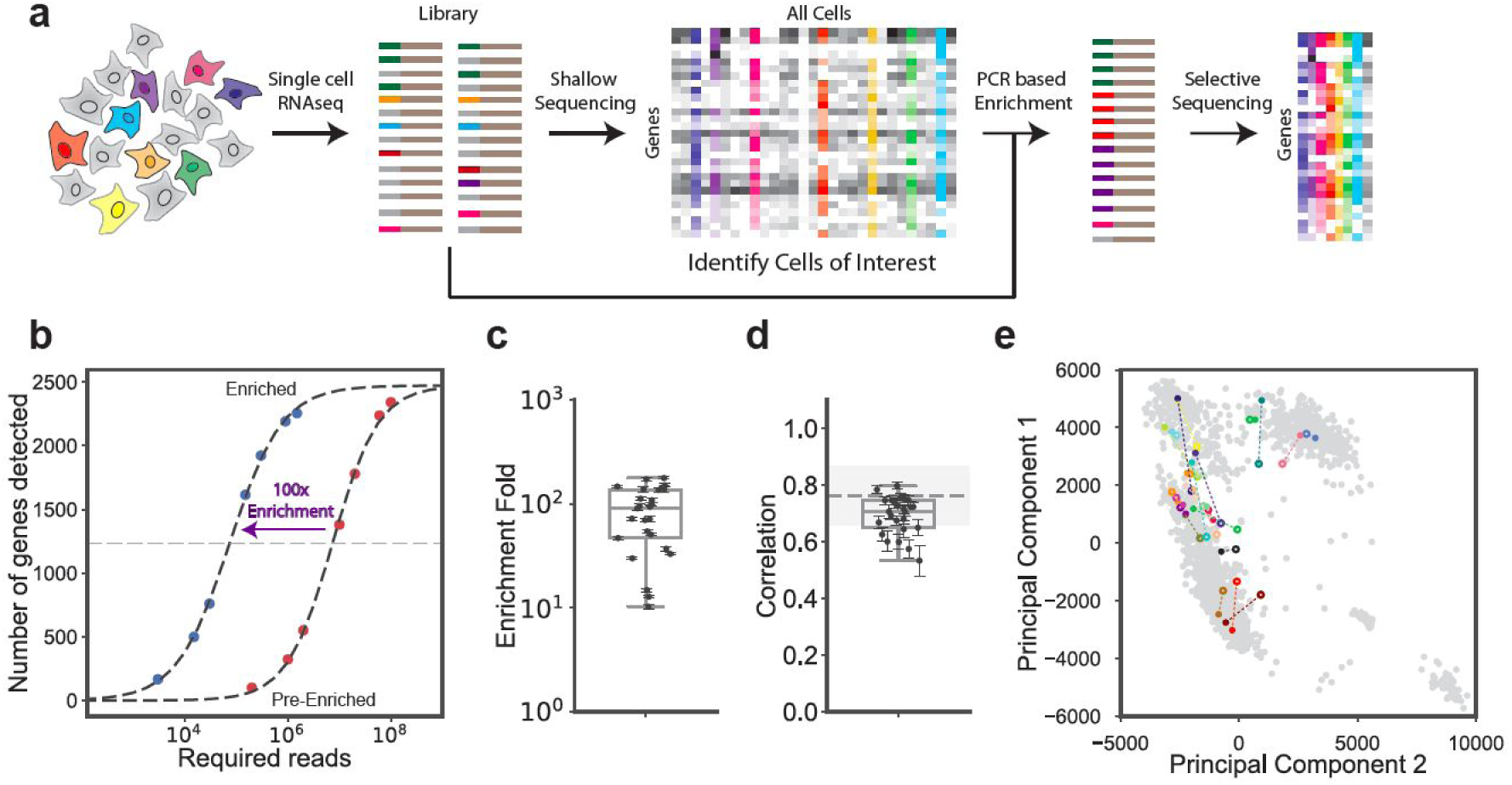
Targeted enrichment of single cells within a pooled RNA-seq sequence library. (a) Workflow showing enrichment based on single cell barcodes on the 5’ end of molecules. Target cells (barcodes) of interest are identified based on shallow sequencing of the original pooled library. PCR with barcode-specific primers is used to create a new sequence library enriched for reads from the target cells. (b) Example enrichment plot for a single target cell from a 10-plex-enrichment reaction. The original library was deeply sequenced as a control to identify gene expression detected in the target cell. Enrichment fold for the cell is read as the fold-difference in overall sequencing effort to detect 50% of the maximum detectable number of genes. (c) Distribution of enrichment-fold values for 24 targeted cells amplified in 10-plex PCR enrichments and (d) the correlation of gene expression profiles pre/post-enrichment for the same 24 cells calculated across genes with at least one detected unique molecular identifier (UMI) in the original sequencing library. The dashed line and shaded region represent the mean and two standard deviation of bootstrap replicates of the 24 cells original gene expression profiles and show the best correlation achievable given the read sampling, UMI sampling, and distribution of expression levels across genes in these specific cells. Error bars represent +/- two SD. (e) Principal component analysis (PCA) of the 24 cells showing the position of the cells based on the expression profiles from the original deep sequenced library (closed circles) and the enriched library (open circles), where each color represents one cell/barcode.

This framework was then applied to target putative AS DCs by combining enrichment by FACS with PCR-based molecular enrichment to target the extremely rare cells. Only 1 million reads were needed to identify some of the key discriminating genes expressed in 9 targeted putative AS DCs captured in the enriched LIN^−^HLA-DR^+^ library. Expression of these AS DC-discriminating genes were either not detectable or resulted in extremely low counts at the same level of sequencing effort in the original, non-enriched library (Supplementary Fig. 7). While the biological role of AS DCs remain to be fully elucidated, the discovery study^7^ reported several properties relevant to the design of new therapeutic and vaccination modalities, highlighting the need to develop strategies to enrich and profile rare cells across from many different samples to decipher their unique properties.

Our results demonstrate that individual cells can be enriched from complex single-cell libraries and that the enriched libraries faithfully represent the targeted cells’ original expression profiles. Selective molecular enrichment of target cells from large pooled single-cell sequencing libraries promises to reduce the sequencing effort required to profile rare cells by one to two orders of magnitude while simultaneously enabling deep sequencing of high-information-content cells.

